# Mosquito population control by aquatic predators: a global meta-analysis of predation efficacy by fish and odonate naiads

**DOI:** 10.64898/2026.08.16.742609

**Authors:** Jesús D. Nuñez, Jolle W. Jolles, Frederic Bartumeus

## Abstract

1. Biological control of mosquitoes using aquatic predators offers a sustainable alternative to chemical insecticides, yet the specific predator and prey functional traits that govern consumption efficacy remain poorly quantified at a global scale.
2. We conducted a global meta-analysis of 755 effect sizes across 59 studies to evaluate how predator identity (fish vs. odonate naiads), body size, dietary guilds, and prey characteristics influence consumption rates. Using multilevel models and robust publication-bias corrections, we quantified predation efficiency, expressed throughout as the consumption rate (CR, larvae predator □¹ h □¹), while accounting for methodological variations across experimental designs. The primary literature itself proved geographically skewed towards Asia (chiefly India), with Africa, the Americas, and Europe markedly under-represented.
3. Grouping predators solely by broad taxonomic identity concealed the central pattern in our data. Although naiads outperformed fish when compared directly within the same studies, this taxon-level difference was driven almost entirely by non-mosquitofish species, the least efficient predator group overall. Mosquitofish (*Gambusia* spp.) and dragonfly naiads were statistically indistinguishable from one another, indicating that dietary specialisation, not taxonomic identity, is the stronger predictor of predation efficacy. Predator body size strongly and positively predicted consumption rates in naiads—driven primarily by dragonflies—but showed no significant or negative relationship in fish.
4. Methodological traits heavily structured the extreme heterogeneity observed across studies; notably, exposure time acted as a severe rate-suppressor, where prolonged assays drastically underestimated per-hour consumption rates due to satiety or handling constraints. Nevertheless, a combined model incorporating all significant ecological moderators simultaneously explained a substantial share of the between-study variance, confirming that predator-prey dynamics in these systems are highly predictable from functional traits.
5. Effective biological control cannot rely on broad taxonomic assumptions but requires evidence-based trait-matching. Management programs should prioritise body size when deploying insect predators, selecting the largest individuals within species known to consume mosquito larvae and favour insectivorous fish species over generalists. Crucially, because short-term laboratory assays artificially inflate efficacy, multi-duration assessments are essential to accurately scale up biocontrol predictions from experimental arenas to complex, real-world ecosystems.

## 1. Introduction

Mosquito-borne diseases and the escalating pressure they exert on global health systems represent one of the most critical humanitarian and ecological challenges of our time (Abbasi, 2025). Malaria alone accounts for approximately 247 million cases and 619,000 deaths annually, while the incidence of arboviral diseases like dengue has surged 30-fold over the past five decades (World Health Organization [WHO], 2023). Furthermore, anthropogenic drivers, including climate change (CITA FALTANTE: Reiter2001), accelerated globalisation, and the range expansion of highly invasive vectors such as *Aedes albopictus* and *A. aegypti* have rendered temperate regions increasingly hospitable to transmission (Kraemer et al., 2019; Wilson et al., 2020). While chemical insecticides have historically formed the backbone of vector management, widespread physiological resistance now severely threatens their long-term viability (Hemingway et al., 2016; Ranson & Lissenden, 2016), besides its strongly pervasive effects by contaminating the environment and harming non-target aquatic organisms (Li et al., 2025; van den Berg et al., 2021). Consequently, there is increasing need for biological control to mitigate vector proliferation without further disrupting the integrity of aquatic communities (Benelli et al., 2026; Alomar, 2025), not just as a sustainable alternative, but as an ecological imperative.

Biological control of mosquito larvae has been practiced globally for over a century, with aquatic predators, particularly fish, being the most widely deployed agents (Benelli et al., 2016; Dye-Braumuller et al., 2020). The mosquitofish (*Gambusia affinis* and *G. holbrooki*) is the emblematic example: intentionally introduced worldwide throughout the twentieth century as a mosquito-control agent (Pyke, 2008), yet simultaneously responsible for large-scale disruption of native aquatic communities (Arthington, 1989; Kurtul et al., 2024; Gil-Luna et al., 2025) and now listed among the world’s worst invasive alien species (Lowe et al., 2000; GISD, 2024). This tension between biocontrol utility and ecological harm underscores the need to move beyond a narrow reliance on introduced species toward native alternatives. Odonate naiads (dragonfly and damselfly larvae) are particularly promising in this respect: they exert powerful regulatory pressure on mosquito larvae within natural wetlands and are widely distributed across biogeographic regions, yet their comparative efficacy relative to fish remains disproportionately understudied (Fan et al., 2025; Priyadarshana & Slade, 2023; Rawani, 2023). More broadly, optimising bio-intervention strategies requires shifting away from broad taxonomic generalisations (e.g., fish vs. naiad) towards a mechanistic focus on the functional traits that actually predict predator performance (Onen et al., 2024). Translating this shift (from asking which taxon consumes more prey to asking which traits predict predation) into an evidence-based framework is far from straightforward.

A first step is to review what the scientific literature already offers. Synthesising this primary literature, however, requires first resolving three critical knowledge gaps. First, a methodological gap: primary studies employ highly variable exposure durations (ranging from minutes to 24 hours), which means raw consumption rates cannot be compared without explicitly accounting for temporal scaling and predator satiety over time (Walshe et al., 2017). Second, a structural gap exists in cross-taxon comparisons; the body size ranges of fish and odonate naiads are largely non-overlapping (typically 15–300 mm vs. 4–45 mm, respectively), meaning that observed differences in predation efficiency between groups risk confounding taxonomic identity with body size (Brose et al., 2006). Third, no study has yet integrated fish and macroinvertebrates within a single unified analytical framework; past datasets for these groups were assembled through disparate search strategies and distinct inclusion criteria, preventing robust cross-taxon comparisons (Nakagawa & Santos, 2012). These gaps are further compounded by the geographic bias of the literature, which relies heavily on laboratory assays from Asia, limiting generalisability to the American, African, and European regions bearing the highest vector burdens (World Health Organization [WHO], 2023; Dambach et al., 2020).

To address these gaps, we present a systematic review and global meta-analysis quantifying the larval consumption rates of aquatic predators against mosquito vectors. Our specific objectives are to: (1) establish a baseline of predation efficiency across fish and odonate naiad predators using a unified metric; (2) examine how key functional traits—specifically dietary guild and body size (Peters, 1983; Brose et al., 2006)—predict larval consumption efficiency across predator groups; (3) determine how prey characteristics (mosquito genus and larval stage) interact with predator identity to shape vulnerability; and (4) characterise the contribution of experimental design variation, particularly exposure duration, to the extreme heterogeneity observed in efficacy estimates. By resolving these methodological and ecological uncertainties, this synthesis aims to advance from ad-hoc predator introductions toward a predictable, trait-based framework for evidence-based vector management.

## 2. Materials and Methods

### 2.1 Literature search and study selection

We conducted a systematic review following the PRISMA-EcoEvo guidelines (O’Dea et al., 2021). We searched Web of Science and Scopus (1 January 1980 to 23 June 2024) to retrieve fish predation datasets using the boolean string: (“fish predation” OR “predatory fish” OR “aquatic predator”) AND (“mosquito larva” OR “larval mosquito”) AND (“biological control” OR “mosquito control” OR “larvivorous fish*”).* This initial search yielded 1,926 unique records after de-duplication using the litsearchr R package (Grames et al., 2019); title and abstract screening was then managed in metagear (Lajeunesse, 2016); the complete screening flowchart detailing the attrition of records can be found in Figure S1. For macroinvertebrate predators, our dataset was initially derived from the comprehensive database compiled by Priyadarshana and Slade (2023), which we supplemented with an updated, targeted search spanning January 2022 to July 2024 using the terms: mosquito AND (dragonfl *OR damselfl* OR odonat*), identifying two additional primary studies (yielding 14 effect sizes).

Fish and naiad effect sizes were identified through independent search strategies — a global database search for the former and the existing compilation by Priyadarshana and Slade (2023) for the latter — and differences in literature sourcing could in principle introduce systematic biases into cross-taxon comparisons. To directly address this concern, we identified six primary studies within our synthesis—hereafter designated as overlap studies (P5, P8, P9, P10, P28, and P29 in Supplementary Material Data S1)—that experimentally tested both fish and odonate naiads simultaneously under identical, standardized environmental conditions. These studies provide a direct, within-study comparison that is entirely independent of between-study methodological differences, and we used this to conduct a pre-specified sensitivity analysis restricted to this overlap subset (see Section 2.7).

To ensure data comparability and ecological relevance, studies were required to meet five strict inclusion criteria. First, they had to experimentally quantify predation as the proportion or absolute number of larvae consumed, allowing for the mathematical derivation of consumption rates. Second, target prey was restricted exclusively to larvae of the genera *Aedes*, *Anopheles*, or *Culex*; these genera were chosen because they represent the primary vectors of global public health concern (transmitting dengue, malaria, and West Nile virus, respectively) and comprise the vast majority of target organisms in management programs. Third, studies had to report sufficient statistical data (sample sizes, means, and variances, or raw counts) for effect size calculation; those lacking this information were excluded at full-text screening due to missing variance metrics or irrecoverable graphical data. Fourth, the predator identity had to be resolved to at least the family level to enable functional trait assignment. Fifth, studies had to be published in peer-reviewed journals rather than grey literature (e.g., technical reports or conference abstracts) to ensure that all synthesized methodologies had undergone rigorous independent evaluation.

### 2.2 Data extraction and standardisation

To extract the data for our meta-analyses, for each paper we used a standardised template to capture thirteen variables organised into six core categories (see Supplementary Table S1): (i) bibliographic and geographic information (publication year, and the country and continent in which the study was conducted, which we used to characterise the taxonomic and geographic coverage of the literature, detailed below); (ii) predator taxonomic identity (species, family, and broad predator group); (iii) predator functional traits (body length and dietary guild); (iv) prey characteristics (mosquito genus and larval instar); (v) experimental design (exposure duration, number of replicates, and enclosure type); and (vi) outcome data (sample size, number of larvae consumed, and the statistics required to derive sampling variance). This structure ensured that every effect size could be traced back to the full set of biological and methodological covariates required for the moderator and sensitivity analyses described below. For studies that reported results exclusively in graphical format, we extracted means and variances using the metaDigitise R package (Pick et al., 2019). When one of the thirteen extracted variables was missing from a primary study, we contacted the corresponding authors up to two times over a four-week period; studies that failed to provide the required variance or sample size metrics after this window were excluded from the synthesis.

To investigate the role of predator dietary specialization in driving biocontrol outcomes, we classified the feeding modes of the studied fish species using a two-step procedure. First, we extracted ecological descriptions directly from the primary studies; when authors explicitly detailed the feeding habits of their experimental subjects, we adopted their classification. When such specific descriptions were lacking, we consulted FishBase (Froese & Pauly, 2025) and its underlying primary literature. This then allowed us to assign species to one of three functional categories: insectivorous (diet consisting predominantly of insects and other arthropods), omnivorous (diet including a mix of plant material, detritus, and macroinvertebrates), and herbivorous (diet consisting predominantly of plant material or algae). To ensure data robustness, species described as strictly piscivorous or with highly specialized feeding mechanisms (e.g., filter-feeders) were excluded from the dietary guild analysis due to insufficient representation in the literature. Odonate naiads were assumed to be obligate generalist macroinvertebrate predators across all taxa considered (Corbet, 1999).

Because exposure times varied substantially across the compiled literature (ranging from almost 5 min to 24 h), raw proportions of consumed larvae would conflate true predation velocity with experimental duration. We therefore standardized all predation outcomes by expressing them as log-natural incidence rates (IRLN), a metric specifically designed to model count data per unit of time under a Poisson approximation (Viechtbauer, 2010). The effect sizes and their corresponding sampling variances (v□) were computed directly from the raw count and exposure data reported in each study using the escalc function (with argument measure = “IRLN”) within the metafor R package (Viechtbauer, 2010), formulated as:

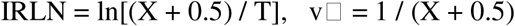

where X represents the number of mosquito larvae consumed, and T = n_pred × t_h denotes the total predator-hours of exposure (calculated as the number of active predators multiplied by the experimental duration in hours). The Haldane correction (+0.5) was applied to both the numerator and denominator to stabilize variances and prevent computational failures in zero-count observations (Viechtbauer, 2010). Crucially, this mathematical framework allowed us to standardize the odonate naiad dataset updated from Priyadarshana and Slade (2023) with our newly compiled dataset with the same temporal metrics. Finally, we computed back-transformed values, which we refer to throughout as the consumption rate (CR = eIRLN), to provide an intuitive ecological interpretation representing the absolute number of larvae consumed per predator type, per individual-hour, where CR = 1 serves as the natural null baseline.

### 2.3 Statistical analysis

#### 2.3.1 Predictors and model structure

To evaluate how different traits modulate larvae predation efficacy, we employed a dual frequentist and Bayesian multilevel meta-regression approach that allowed us to simultaneously estimate the effects of multiple predator, prey, and methodological moderators while accounting for non-independence among effect sizes drawn from the same study. Our response variable across all models was the log-incidence rate (IRLN) described above, which explicitly standardises predation outcomes for exposure duration and the number of predators used. We examined six moderators a priori: (1) predator identity (fish vs. naiads; and at finer resolution: mosquitofish, non-mosquitofish, dragonfly, damselfly); (2) mosquito genus (*Aedes*, *Anopheles*, *Culex*); (3) larval stage (early: instars I–II; late: instars III–IV); (4) predator body size (mm, continuous); (5) fish feeding mode (insectivorous, omnivorous, herbivorous); and (6) number of experimental replicates [log-transformed, log(repl)] (Nakagawa & Santos, 2012; Nakagawa et al., 2015). Two-way interactions were tested between predator type and each prey moderator. Body size was z-standardised using the global mean and SD of all observations with recorded size (x = 38.9 mm, SD = 47.8 mm, n = 334). Given that fish and naiads occupy largely non-overlapping size ranges (fish: 15–300 mm; naiads: 4–45 mm), body size was analysed separately within each broad predator group. Within-group slopes (SD _naiad_ = 11.4 mm; SD _fish_ = 61.7 mm) are reported in Supplementary Table S10. Exposure time was modelled as a continuous covariate (raw hours).

#### 2.3.2 Variance partitioning and heterogeneity

To quantify total variance explained by measured moderators, following a pseudo-R² approach analogous to that of Nakagawa and Schielzeth (2013), we fitted a global model including all six moderators simultaneously and calculated the proportional reduction in variance components relative to the intercept-only model. We did this separately for the between-study (σ²□) and within-study (σ²□) variance components, so that the relative contribution of the fixed effects to each level of the random-effects structure could be distinguished. All models shared a two-level hierarchical random-effects structure: a random intercept for study identity (StudyID) and one for effect size identity (SubID) nested within studies, specified in metafor as list(∼1|StudyID,∼1|SubID), estimating separate between-study (σ²□) and within-study (σ²□) variance components. Frequentist analyses used rma.mv (restricted maximum likelihood; REML; Viechtbauer (2010)). Bayesian analyses used brms (Bürkner, 2017) with Normal(0,1) priors for regression coefficients and half-t(3) for variance components, run with four MCMC chains (6000 iterations, 2000 warm-up). Convergence was assessed via R□ < 1.01 (Vehtari et al., 2021) and ESS > 400 (Vats & Knudson, 2021) (full convergence diagnostics and Bayesian–frequentist comparison in Supplementary Table S2). Heterogeneity in predation efficacy across studies was quantified using multilevel I² (Nakagawa et al., 2015), which partitions total heterogeneity into within-study and between-study components appropriate for our two-level random-effects structure, via dmetar (Harrer et al., 2019).

Publication bias was assessed with four complementary tests, each sensitive to a different signature of small-study or selective-reporting bias: Kendall’s rank-correlation (Begg & Mazumdar, 1994), which tests for an association between effect size and its standard error; Egger’s regression adapted for multilevel models (Egger et al., 1997; Rodgers & Pustejovsky, 2020), which tests for funnel-plot asymmetry; trim-and-fill (Duval & Tweedie, 2000), which imputes hypothetically missing studies to restore funnel-plot symmetry; and PET-PEESE (Precision-Effect Test and Precision-Effect Estimate with Standard Error; (Stanley & Doucouliagos, 2014)), a meta-regression approach that models effect size as a function of its standard error (PET) or variance (PEESE) to estimate the effect size expected at infinite precision. Because trim-and-fill performs poorly under the extreme heterogeneity typical of ecological meta-analyses (Terrin et al., 2003), we treat PET-PEESE as the more reliable correction where the two disagree (full results in Supplementary Table S4). Sensitivity analyses included leave-one-out analyses (Supplementary Table S5), Cook’s distance (> 4/k threshold) and studentized deleted residuals (|z| > 3) for outlier diagnostics, and cluster-robust standard errors (type CR2; Rodgers and Pustejovsky (2020)) to verify significance of main effects (Supplementary Table S6). Across all models, these diagnostics indicated good robustness: publication-bias corrections, leave-one-out analysis, and outlier diagnostics showed no undue influence of individual studies, and Bayesian estimates were concordant with frequentist results throughout. Full coefficients for the predator-by-genus, predator-by-stage, and specific-predator-by-stage interaction models are reported in Supplementary Tables S7, S8, and S9 respectively; within-group body-size slopes and the practical, duration-dependent consequences of the exposure-time effect for each predator group are reported in Supplementary Tables S10 and S11. All analyses were conducted in R 4.3.1 (R Core Team, 2023).

## 3. Results

The resulting dataset, comprising the 59 studies included in this meta-analysis in total, was geographically and taxonomically uneven. Of these, 39 (67%) were conducted in Asia, with India alone accounting for 28 studies; Africa and the Americas each contributed fewer than 15% of studies, and Europe was critically under-represented. This imbalance is unlikely to reflect regional research need and more plausibly reflects disparities in research infrastructure and funding; because most laboratory assays used locally sourced predator populations, it also means that population-and strain-level variation in consumption rate outside Asia remains comparatively untested, a limitation we return to in the Discussion. Odonate naiads were represented in 33 unique studies (dragonfly: k = 418; damselfly: k = 81); fish predators appeared in 32 unique studies (k = 256), of which mosquitofish accounted for 12 studies (k = 60).

Across all 755 effect sizes tested, aquatic predators consumed an average of 2.62 larvae per predator per hour (95% CI: [1.87, 3.68]; p < 0.001; Table 1); throughout this synthesis we refer to this back-transformed metric as the consumption rate (CR), following the definition given in Section 2.

**Table 1.**
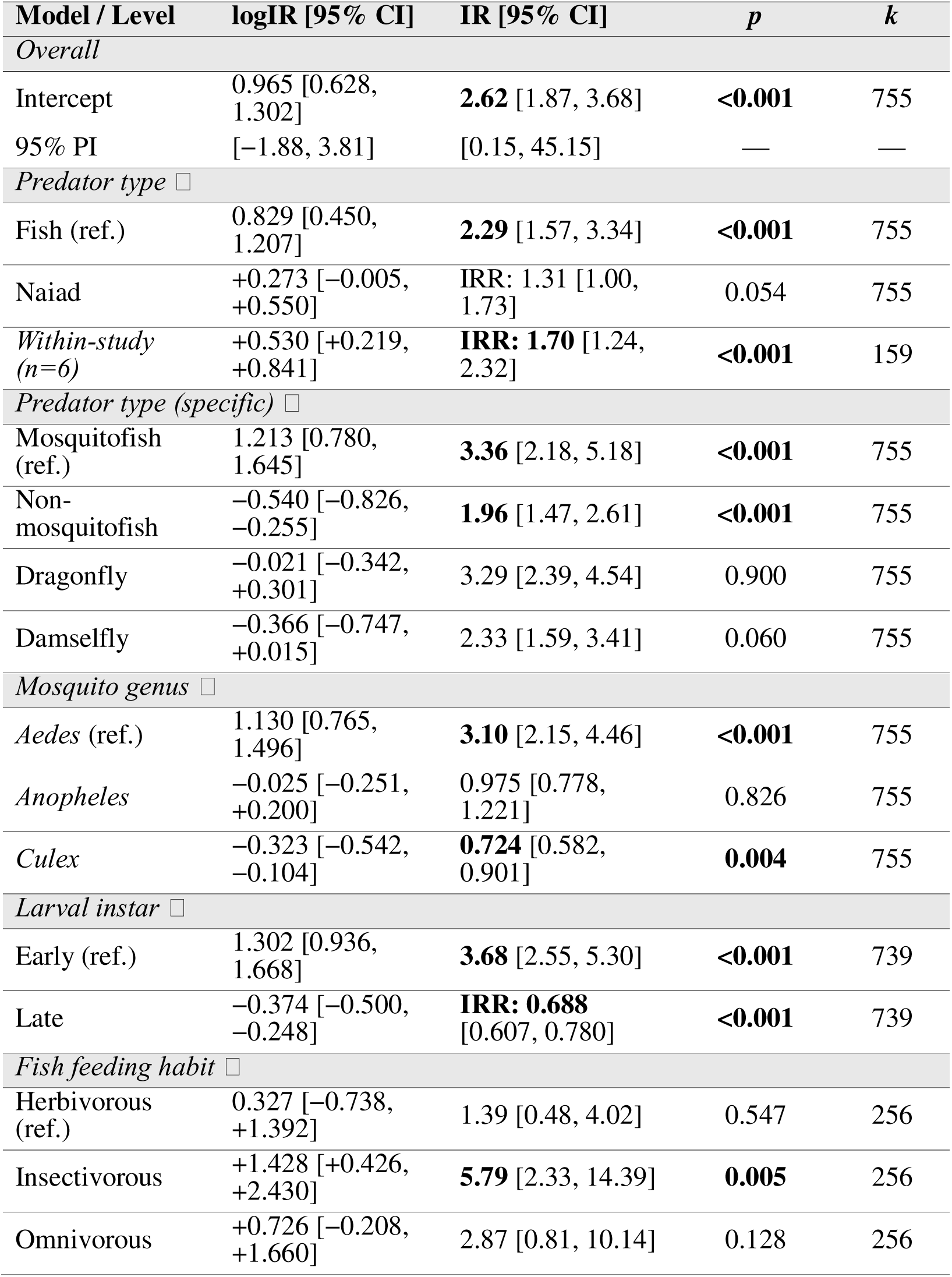
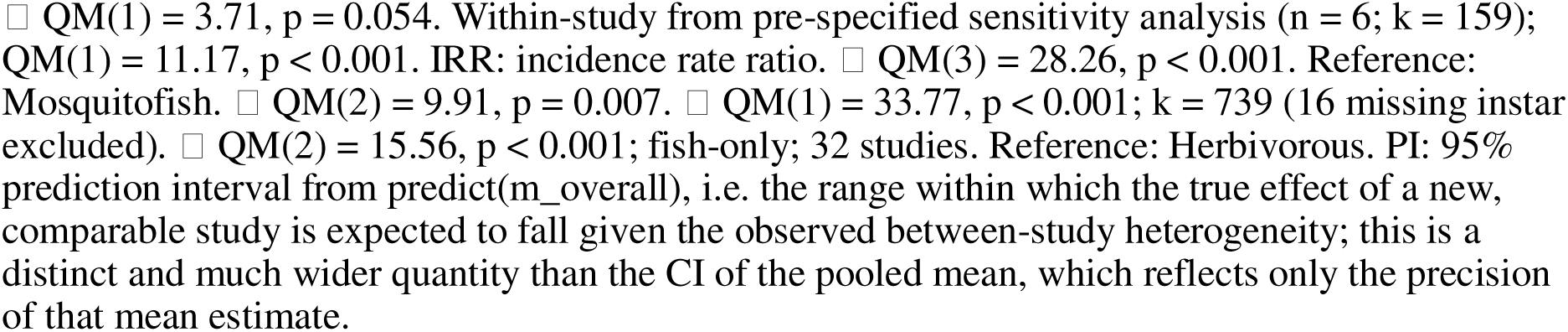
Frequentist meta-analytic models (log incidence rate; rma.mv, REML). IR = elogIR: larvae consumed per predator per hour; IR > 1 indicates predation above the reference unit. The intercept row gives the absolute IR for the reference level; all other rows are contrasts. k: effect sizes. QM: Wald omnibus test (df in parentheses). Bold: p ≤ 0.05. Random effects: list(∼1|StudyID, ∼1|SubID), restricted maximum likelihood (REML); effect size k = 755 from 59 studies except where noted.

| Model / Level | logIR [95% CI] | IR [95% CI] | *p* | *k* |
| --- | --- | --- | --- | --- |
| *Overall* |  |  |  |  |
| Intercept | 0.965 [0.628, 1.302] | **2.62** [1.87, 3.68] | **<0.001** | 755 |
| 95% PI | [-1.88, 3.81] | [0.15, 45.15] | — | — |
| *Predator type* □ |  |  |  |  |
| Fish (ref.) | 0.829 [0.450, 1.207] | **2.29** [1.57, 3.34] | **<0.001** | 755 |
| Naiad | +0.273 [-0.005, +0.550] | IRR: 1.31 [1.00, 1.73] | 0.054 | 755 |
| *Within-study (n=6)* | +0.530 [+0.219, +0.841] | **IRR: 1.70** [1.24, 2.32] | **<0.001** | 159 |
| *Predator type (specific)* □ |  |  |  |  |
| Mosquitofish (ref.) | 1.213 [0.780, 1.645] | **3.36** [2.18, 5.18] | **<0.001** | 755 |
| Non-mosquitofish | -0.540 [-0.826, -0.255] | **1.96** [1.47, 2.61] | **<0.001** | 755 |
| Dragonfly | -0.021 [-0.342, +0.301] | 3.29 [2.39, 4.54] | 0.900 | 755 |
| Damselfly | -0.366 [-0.747, +0.015] | 2.33 [1.59, 3.41] | 0.060 | 755 |
| *Mosquito genus* □ |  |  |  |  |
| *Aedes* (ref.) | 1.130 [0.765, 1.496] | **3.10** [2.15, 4.46] | **<0.001** | 755 |
| *Anopheles* | -0.025 [-0.251, +0.200] | 0.975 [0.778, 1.221] | 0.826 | 755 |
| *Culex* | -0.323 [-0.542, -0.104] | **0.724** [0.582, 0.901] | **0.004** | 755 |
| *Larval instar* □ |  |  |  |  |
| Early (ref.) | 1.302 [0.936, 1.668] | **3.68** [2.55, 5.30] | **<0.001** | 739 |
| Late | -0.374 [-0.500, -0.248] | **IRR: 0.688** [0.607, 0.780] | **<0.001** | 739 |
| *Fish feeding habit* □ |  |  |  |  |
| Herbivorous (ref.) | 0.327 [-0.738, +1.392] | 1.39 [0.48, 4.02] | 0.547 | 256 |
| Insectivorous | +1.428 [+0.426, +2.430] | **5.79** [2.33, 14.39] | **0.005** | 256 |
| Omnivorous | +0.726 [-0.208, +1.660] | 2.87 [0.81, 10.14] | 0.128 | 256 |

□ $QM(1) = 3.71$ , $p = 0.054$ . Within-study from pre-specified sensitivity analysis ( $n = 6$ ; $k = 159$ ); $QM(1) = 11.17$ , $p < 0.001$ . IRR: incidence rate ratio. □ $QM(3) = 28.26$ , $p < 0.001$ . Reference: Mosquitofish. □ $QM(2) = 9.91$ , $p = 0.007$ . □ $QM(1) = 33.77$ , $p < 0.001$ ; $k = 739$ (16 missing instar excluded). □ $QM(2) = 15.56$ , $p < 0.001$ ; fish-only; 32 studies. Reference: Herbivorous. PI: 95% prediction interval from $\text{predict}(m\_overall)$ , i.e. the range within which the true effect of a new, comparable study is expected to fall given the observed between-study heterogeneity; this is a distinct and much wider quantity than the CI of the pooled mean, which reflects only the precision of that mean estimate.

The meta-analysis revealed extreme heterogeneity in CR across studies (multilevel I² = 99.1%; full variance-partitioning statistics in Supplementary Table S3). Robustness diagnostics (Section 2.3) confirmed this heterogeneity was not attributable to publication bias or influential studies: we treat PET-PEESE (Precision-Effect Test and Precision-Effect Estimate with Standard Error; (Stanley & Doucouliagos, 2014)), a meta-regression approach that regresses effect size on its standard error (PET) or, where a genuine effect is indicated, on its variance instead (PEESE), to estimate the effect size expected at infinite precision (i.e. zero standard error). Because trim-and-fill performs poorly under the extreme heterogeneity typical of ecological meta-analyses (Terrin et al., 2003), we treat PET-PEESE as the more reliable correction where the two disagree (full results in Supplementary Table S4), cluster-robust standard errors confirmed all main effects (Supplementary Table S6), and no single study unduly influenced the overall estimate (leave-one-out diagnostics in Supplementary Table S5). Bayesian models converged well and were concordant with the frequentist results across all moderators, including for the sparsely represented herbivorous-feeding and late-instar categories (full posterior estimates in Supplementary Table S2).

### 3.1 Exposure time

Exposure time was a significant negative predictor of CR across all groups (β = −0.072, 95% CI: [−0.080, −0.063], p < 0.001, R² = 24.2%; Table 2): a 24-hour trial yielded an estimated marginal mean only 19% of that of a 1-hour trial. The effect was strongest in non-mosquitofish fish (β = −0.109/h; 12-fold overestimate) and weakest in damselfly naiads (β = −0.038/h; Table 2; the practical, duration-dependent consequences of these slopes for each predator group are tabulated in Supplementary Table S11). A fraction of this negative slope is mathematically inherent to the underlying incidence-rate metric; disentangling true biological prey depletion or predator satiation from this mathematical component requires factorial designs that orthogonally vary exposure time and prey density.

**Table 2.** Meta-regression slopes for predator body size (z-scored; global 1 SD = 47.8 mm) and exposure time (hours) on log incidence rate (IRLN). Body size slopes from separate within-group models. Within-group standardised slopes in Supplementary Table S10. Bold: p < 0.05.

| Moderator | Group | $\beta$ | 95% CI | $p$ | Sig. |
| --- | --- | --- | --- | --- | --- |
| *Body size (global z-score; 1 SD = 47.8 mm)* |  |  |  |  |  |
|  | Overall | +0.122 | [−0.029, +0.272] | 0.113 | ns |
| Broad | Naiad | **+0.693** | **[+0.249, +1.137]** | **0.002** | **\*\*** |
|  | Fish | −0.155 | [−0.391, +0.082] | 0.200 | ns |
| Specific | Dragonfly | **+0.579** | **[+0.171, +0.987]** | **0.006** | **\*\*** |
|  | Damselfly † | +3.449 | [+0.388, +6.509] | 0.027 | * |
|  | Mosquitofish | +0.143 | [−1.395, +1.682] | 0.855 | ns |
|  | Non-mosquitofish | −0.224 | [−0.473, +0.025] | 0.078 | . |
| *Exposure time (h, raw scale)* |  |  |  |  |  |
|  | Overall | **−0.072** | **[−0.080, −0.063]** | **<0.001** | **\*\*\*** |
| Broad | Naiad | **−0.050** | **[−0.057, −0.042]** | **<0.001** | **\*\*\*** |
|  | Fish | **−0.102** | **[−0.121, −0.084]** | **<0.001** | **\*\*\*** |
| Specific | Dragonfly | **−0.053** | **[−0.061, −0.045]** | **<0.001** | **\*\*\*** |
|  | Damselfly | **−0.038** | **[−0.056, −0.020]** | **<0.001** | **\*\*\*** |
|  | Mosquitofish | **−0.057** | **[−0.094, −0.020]** | **0.002** | **\*\*** |
|  | Non-mosquitofish | **−0.109** | **[−0.131, −0.088]** | **<0.001** | **\*\*\*** |

† Damselfly slope implies a 31-fold per-hour increase per SD; biologically implausible ( $n = 34$ , range 4–25 mm).
\*\*\* $p < 0.001$ ; \*\* $p < 0.01$ ; \* $p < 0.05$ ; . $p < 0.10$ ; ns $p \geq 0.10$ .

**Table 3.** Evidence-based practical recommendations for trait-based biocontrol agent selection. IR: expected larvae predator □¹ h □¹ for the recommended configuration. Overestimation factor: the factor by which a 1-hour assay overestimates the sustained, 24-hour per-hour rate for that predator group (e.g. a value of 3× means the 1-hour estimate is three times higher than the 24-hour estimate), derived from the exposure-time slopes in Supplementary Table S11.

| Predator | Optimal target | Body size | Min. assay | Expected IR | 1h/24 h |
| --- | --- | --- | --- | --- | --- |
| Dragonfly naiad | *Aedes* (all stages) | Maximise (>20 mm) | ≤6 h | 3–10 | 3× |
| Mosquitofish | *Aedes* early instars | Not predictive | ≤2 h | 3–4 | 4× |
| Insectivorous fish | Any genus | Not predictive | ≤6 h | 4–6 | 10× |
| Damselfly naiad | *Aedes/Anoph.* | Larger if available | ≤6 h | 2–3 | 2× |
| Non-mosquitofish | *Aedes/Anoph.*; avoid *Culex* late | Not predictive | ≤6 h | <2 | 12× |
| *Avoid: any predator vs. Culex late instars in 24-h assays (estimated IR ≈ 0.08).* |  |  |  |  |  |

### 3.2 Taxonomic and ontogenetic patterns in predation

The central pattern in our data is not simply that naiads outperform fish, but that predator identity at a finer taxonomic resolution matters far more than the broad fish-versus-naiad distinction. At the coarse taxonomic level, naiads showed a trend towards higher CR than fish (QM(1) = 3.71, p = 0.054; QM: Wald-type global test statistic for the moderator, chi-squared-distributed with the degrees of freedom given in parentheses; Fig. 2b; 3.00 vs. 2.29 larvae predator □¹ h □¹ (i.e. CR), respectively; incidence rate ratio [naiad:fish] = 1.31, 95% CI: [1.00, 1.73]), a trend corroborated by the within-study sensitivity analysis restricted to the six studies that tested both fish and naiad predators under identical conditions, which confirmed naiads as significantly more efficient than fish specifically within this directly-compared subset (k = 159; incidence rate ratio = 1.70, 95% CI: [1.24, 2.32], QM(1) = 11.17, p < 0.001). However, a finer analysis distinguishing four predator groups (mosquitofish, non-mosquitofish fish, dragonfly naiads, and damselfly naiads — as opposed to the broad, two-level predator type of fish vs. naiads used above) revealed significant heterogeneity among them (QM(3) = 28.26, p < 0.001; Fig. 2c) and a more nuanced pattern: mosquitofish showed the highest CR (3.36, 95% CI: [2.18, 5.18]) and non-mosquitofish fish the lowest (1.96, 95% CI: [1.47, 2.61]; p < 0.001 vs. mosquitofish), while dragonfly naiads (3.29, 95% CI: [2.39, 4.54]; p = 0.900 vs. mosquitofish) and damselfly naiads (2.33, 95% CI: [1.59, 3.41]; p = 0.060 vs. mosquitofish) fell at intermediate-to-high rates. Mosquitofish and dragonfly naiads did not differ significantly from one another (i.e. p = 0.900), and their point estimates were nearly identical (3.29 vs. 3.36); damselfly naiads likewise did not differ significantly from mosquitofish (p = 0.060), although their point estimate was descriptively lower and this comparison approached significance, and together the two naiad groups and mosquitofish clearly outperformed non-mosquitofish fish.

**Figure 1.**
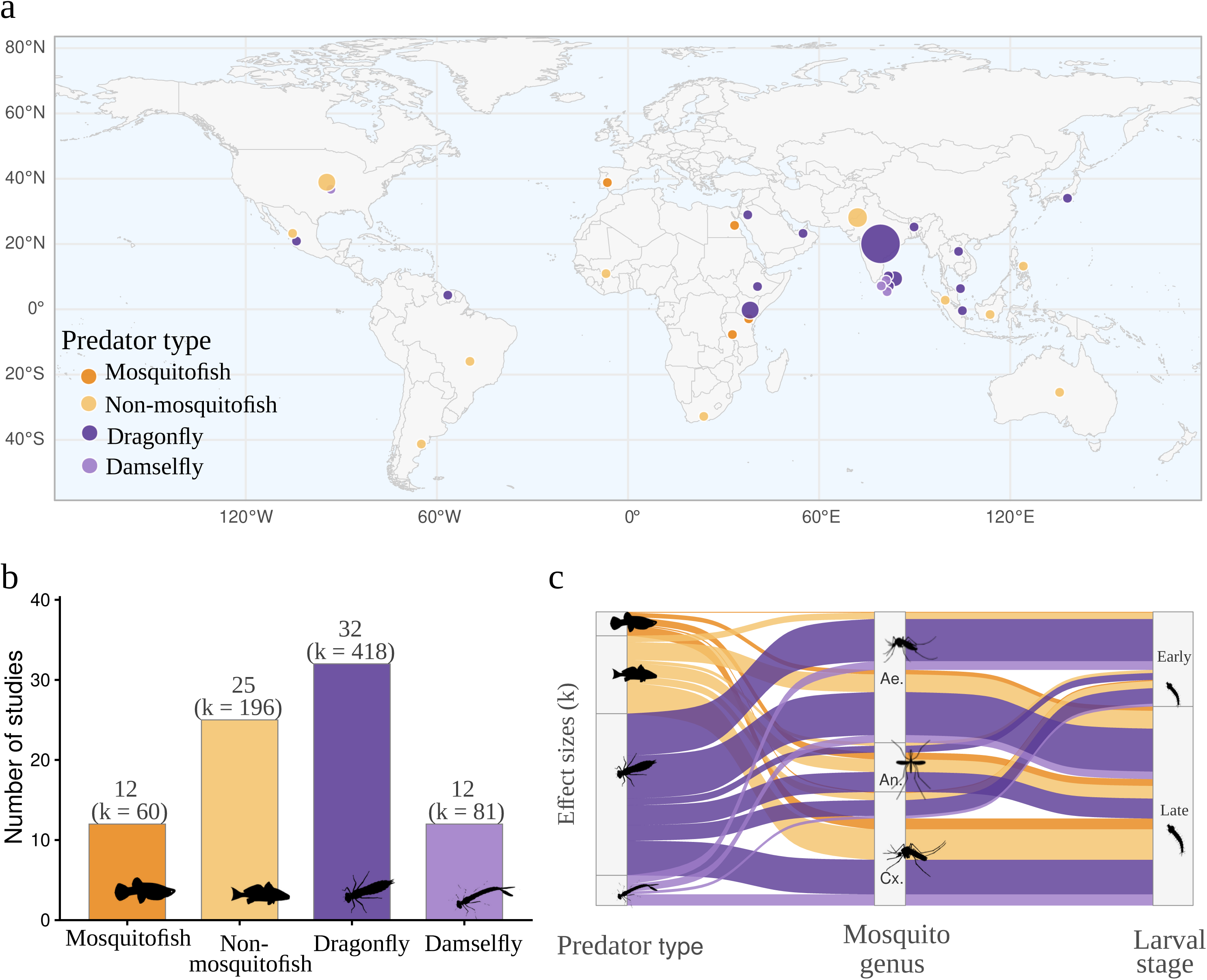
Overview of studies included in the meta-analysis. (a) Geographic distribution; circle size proportional to number of studies per country (range: 1–28; India n = 28); fill colour indicates dominant predator type. (b) Number of studies (bar height) and effect sizes k (parentheses) per predator type: mosquitofish, non-mosquitofish, dragonfly naiads, and damselfly naiads. (c) Sankey diagram linking predator type to mosquito genus (Ae. = *Aedes*; An. = *Anopheles*; Cx. = *Culex*) and larval stage. Icons from PhyloPic.org (CC0 1.0).

**Figure 2.**
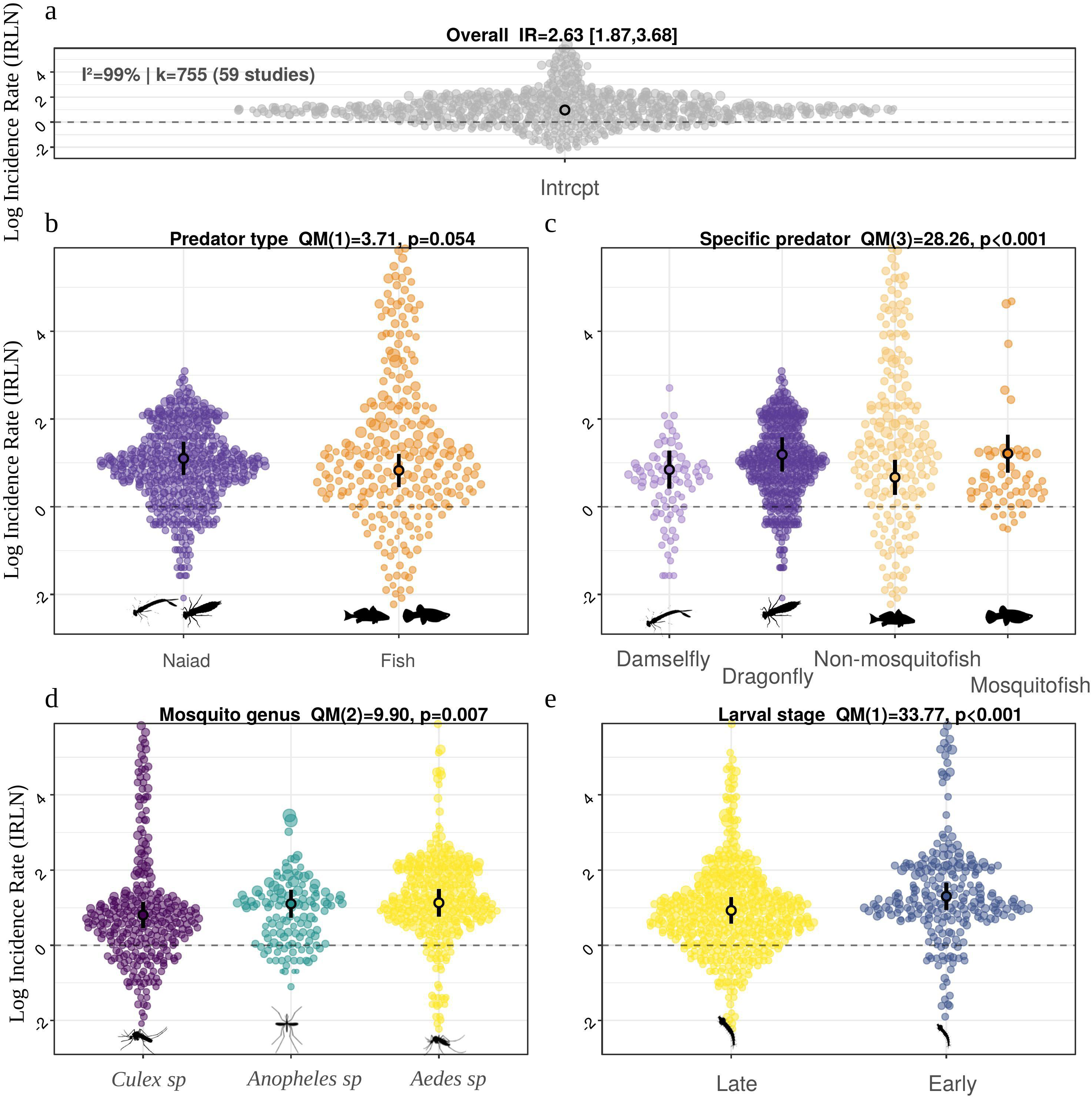
Predation efficiency (IRLN: natural log of larvae consumed per predator per hour) by aquatic predators. (a) Overall mean across all 755 effect sizes (I² = 99.1%). (b) By broad predator type (Fish vs. Naiad); overall difference marginally non-significant (p = 0.054); but see the within-study comparison in Table 1 (six overlap studies) that confirms naiads as significantly more efficient (IRR = 1.70, p < 0.001). (c) By specific predator group (Mosquitofish, Non-mosquitofish, Dragonfly, Damselfly). (d) By mosquito genus. (e) By larval stage. Points: individual effect sizes; horizontal bars and shaded regions: means and 95% CIs. Icons from PhyloPic.org (CC0 1.0).

Developmental stage was a strong predictor of CR: early instars were consumed at higher rates (3.68, 95% CI: [2.55, 5.30]) than late instars (2.53, 95% CI: [1.83, 3.50]), a 31% reduction (QM(1) = 33.77, p < 0.001; Fig. 2e). This general effect was significantly modulated by predator type: at the broad predator-type level, fish showed a steeper 42% decline from early (4.36, [2.88, 6.61]) to late instars (2.52; derived group estimate from the interaction model, full coefficients and CIs in Supplementary Table S8) than naiads, which declined by a shallower 28% (early instars: 3.49, 95% CI: [2.32, 5.24]; late instars: 2.50, 95% CI: [1.80, 3.48]; predator-type × developmental-stage interaction (full coefficients in Supplementary Table S8): p = 0.195, i.e. not statistically significant). Testing this same stage effect within the finer, four-way predator-group classification revealed a significant predator-group × developmental-stage interaction (QM(3) = 35.82, p < 0.001; individual-group effect: QM(7) = 63.74, p < 0.001; full coefficients in Supplementary Table S9; Table 1; Fig. 3b,d). We highlight the two extremes of this four-way pattern here, and report the full set of group-level estimates in Table 1: dragonfly naiads showed the shallowest, and only marginally significant, stage-specific decline (p = 0.019; early: 3.70; late: 2.73; 26% decline), whereas mosquitofish showed by far the steepest (early: 13.65; late: 3.30; 76% decline). Note that the very high early-instar CR for mosquitofish reflects predominantly short-duration (≤ 2 h) assays in this subset, consistent with the exposure-time effect described above (i.e. the shorter the assay, the higher the estimated per-hour CR, as expected if consumption slows over the course of longer trials).

**Figure 3.**
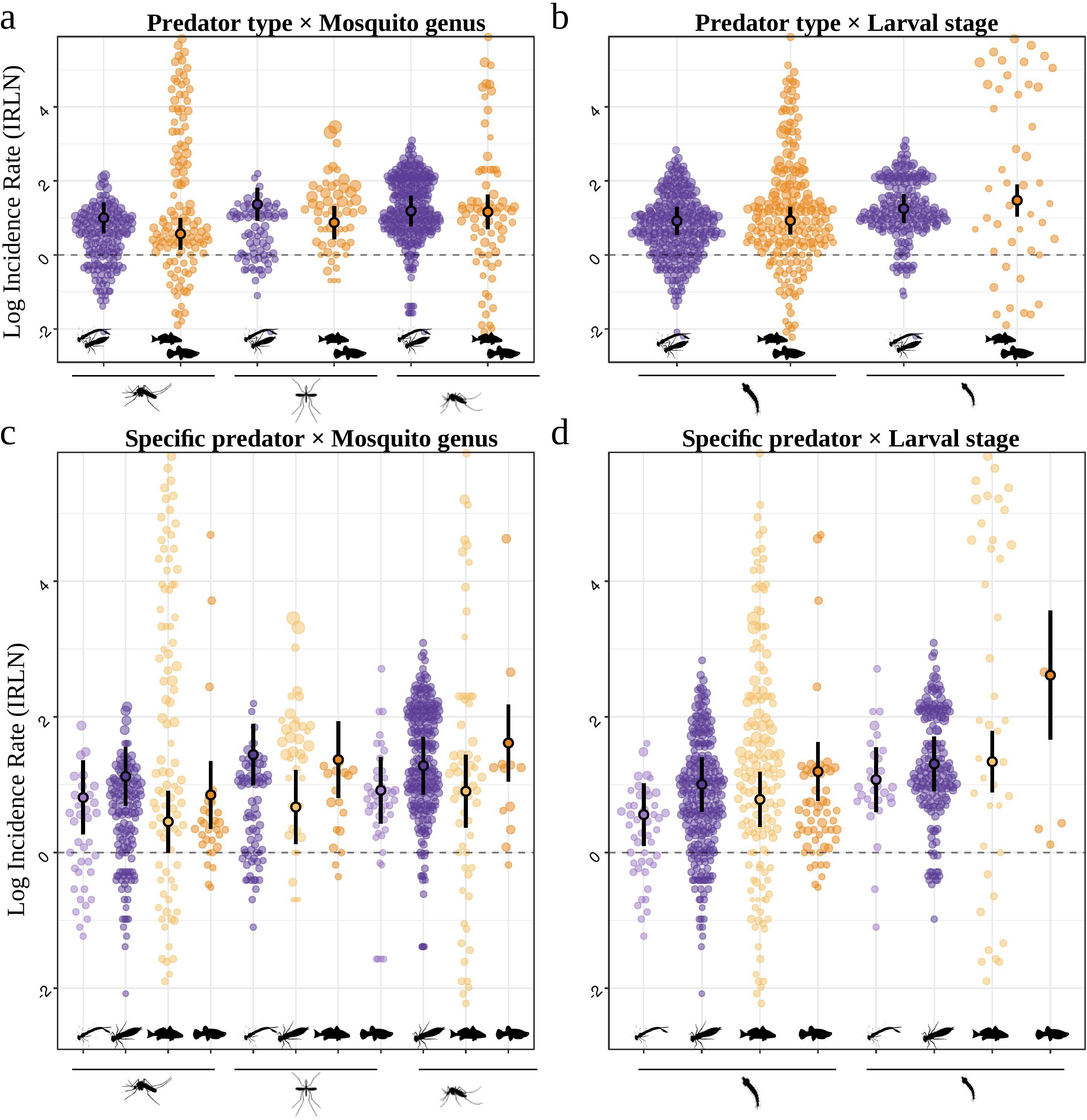
Predation efficiency (IRLN) stratified by mosquito genus and larval instar. (a) Broad predator type × mosquito genus (QM(5) = 17.88, p = 0.003). (b) Broad predator type × larval stage (QM(3) = 35.82, p < 0.001). (c) Specific predator × mosquito genus (Damselfly × Anopheles absent from dataset). (d) Specific predator × larval stage (QM(7) = 63.74, p < 0.001). Icons from PhyloPic.org (CC0 1.0).

Mosquito genus also predicted CR, with *Aedes* predated at the highest rate (3.10, [2.15, 4.46]), followed by *Anopheles* (3.02; p = 0.826 vs. *Aedes*) and *Culex* (2.24; p = 0.004 vs. *Aedes*; overall genus effect: QM(2) = 9.91, p = 0.007; Fig. 2d, Fig. 3a). This general effect was further modulated by predator type (predator-type × mosquito-genus interaction: QM(5) = 17.88, p = 0.003; full coefficients in Supplementary Table S7; Fig. 3a): fish were most efficient against *Aedes* but significantly less so against *Culex* (p = 0.004), whereas naiads showed consistent rates across all three genera (Table 1; Fig. 3a).

Together, these taxonomic and ontogenetic patterns account for much of the extreme heterogeneity noted above: despite an overall I² = 99.1%, a global model (as defined in Section 2.3: a single model with all six moderators included simultaneously) including all six measured moderators simultaneously explained 47.2% of between-study and 29.1% of within-study variance, and the 95% prediction interval (CR = [0.15, 45.15]) that appears uninformative in isolation is largely resolved once these traits are taken into account, confirming that the heterogeneity is structured predominantly by measurable ecological traits rather than random contextual noise.

### 3.3 Predator dietary specialisation

Among fish studies with classifiable feeding mode (k = 256 effect sizes across 32 studies), dietary specialisation significantly predicted CR. Because only two studies used herbivorous fish, we excluded this guild from the formal moderator model and instead compared it descriptively; the formal model therefore tested insectivorous against omnivorous fish, which differed significantly (QM(1) = 15.56, p < 0.001; Table 1; Fig. 4): insectivorous fish showed dramatically higher CR than omnivorous fish (5.79, 95% CI: [2.33, 14.39] vs. 2.87, [0.81, 10.14]). The two herbivorous-fish studies (descriptive only, not modelled) showed a CR of 1.39 [0.48, 4.02], numerically the lowest of the three guilds, consistent with, though not a formal statistical test of, an insectivore-to-herbivore efficacy gradient. The distribution of consumption-rate values for insectivorous fish specifically (not confounded with the broader fish vs. naiad comparison addressed above) showed a bimodal pattern (Fig. 4), with clusters of low-efficacy and high-efficacy studies. This result suggests that further analyses may be needed in order to identify potential drivers (e.g. prey larval stage, arena volume) explaining the differences between these two clusters.

**Figure 4.**
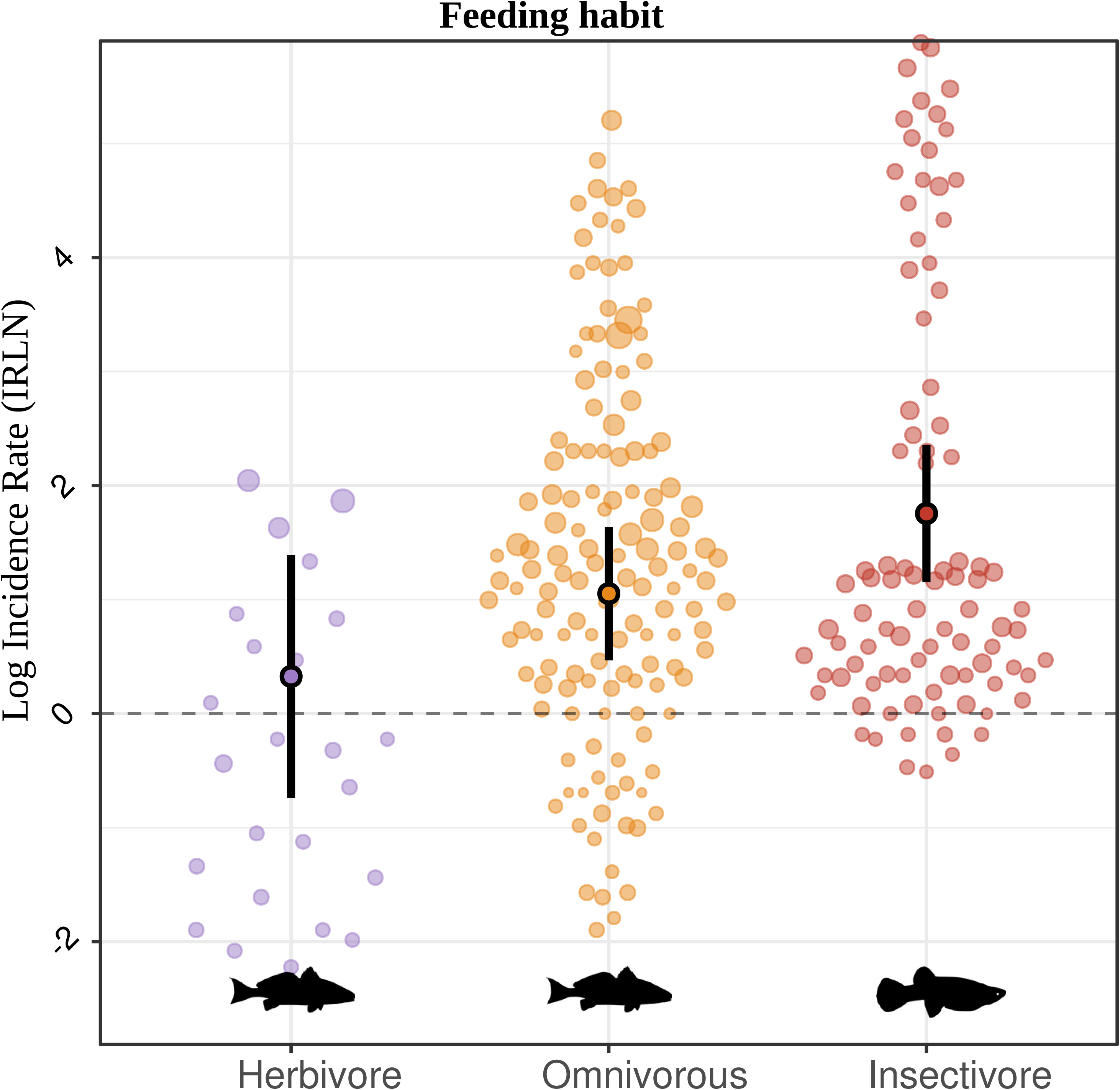
Predation efficiency (IRLN) by fish dietary guild. Herbivorous (k = 24; 2 studies), omnivorous (k = 141; 21 studies), and insectivorous (k = 91; 18 studies) fish. Note the bimodal distribution for insectivores. QM(2) = 15.56, p < 0.001. Points sized by precision (1/√v, where v is the sampling variance of each effect size, as defined in Section 2.2). Icons from PhyloPic.org (CC0 1.0).

### 3.4 Predator body size

Considering the size of the predator, when all data were combined, body size did not have a significant effect on CR (β = +0.122, 95% CI: [−0.029, +0.272], p = 0.113; Fig. 5a). Looking at the two predator types separately, however, revealed that body size did not play a role in predation by fish (β = −0.155, 95% CI: [−0.391, +0.082], p = 0.200; Fig. 5b,c), whereas it was a significant positive predictor among naiads (β = +0.693, 95% CI: [+0.249, +1.137], p = 0.002; Fig. 5d,e), with larger individuals consuming more larvae. Among naiads, the effect was largely driven by dragonfly larvae (β = +0.579, [+0.171, +0.987], p = 0.006; Fig. 5d); damselfly slopes were large but highly uncertain (β = +3.449, 95% CI: [+0.388, +6.509], p = 0.027, n = 34; Table 2; Fig. 5e) and likely unstable given the small sample size underlying this estimate (n = 34). On a biologically interpretable scale, each within-naiad SD increase in body length (11.4 mm) corresponded to an 18% increase in CR (β within = +0.165, p = 0.002; Supplementary Table S10) for mosquito larvae specifically; see the coefficient plot (Fig. 6) for a summary of this contrast across all groups.

**Figure 5.**
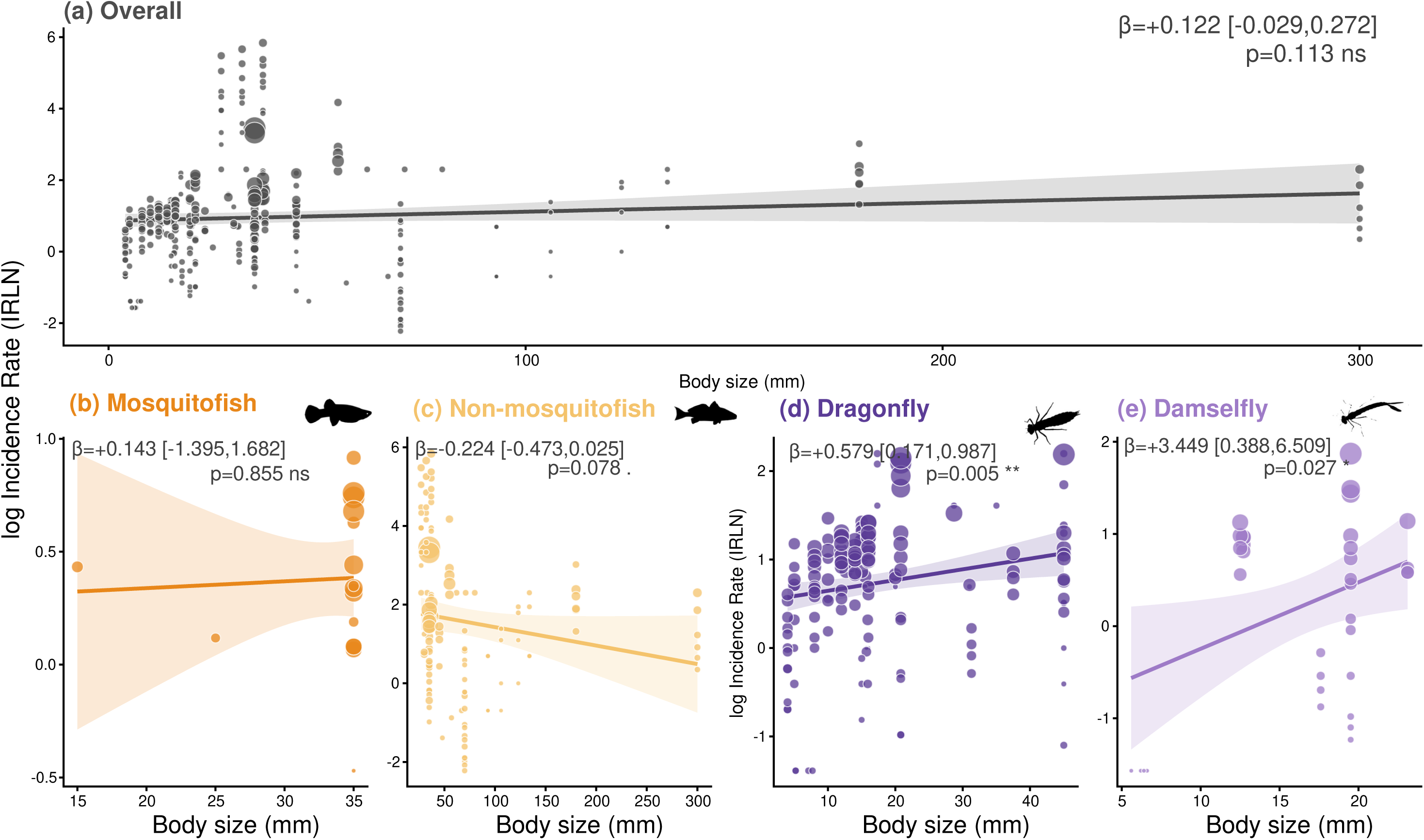
Relationship between predator body size and IRLN. Points scaled by precision (1/√v); lines with 95% CI bands estimated from separate meta-regression models fitted independently within each predator group, rather than a single combined model, reflecting the non-overlapping body-size ranges of fish and naiads (Table 2). (a) Overall (β = +0.122, ns). (b) Mosquitofish (β = +0.143, ns). (c) Non-mosquitofish (β = −0.224, p = 0.078). (d) Dragonfly naiads (β = +0.579, p = 0.006). (e) Damselfly naiads (β = +3.449, p = 0.027; see text for caveats on n = 34). Note contrasting directions between fish (panels b–c) and naiads (panels d–e).

**Figure 6.**
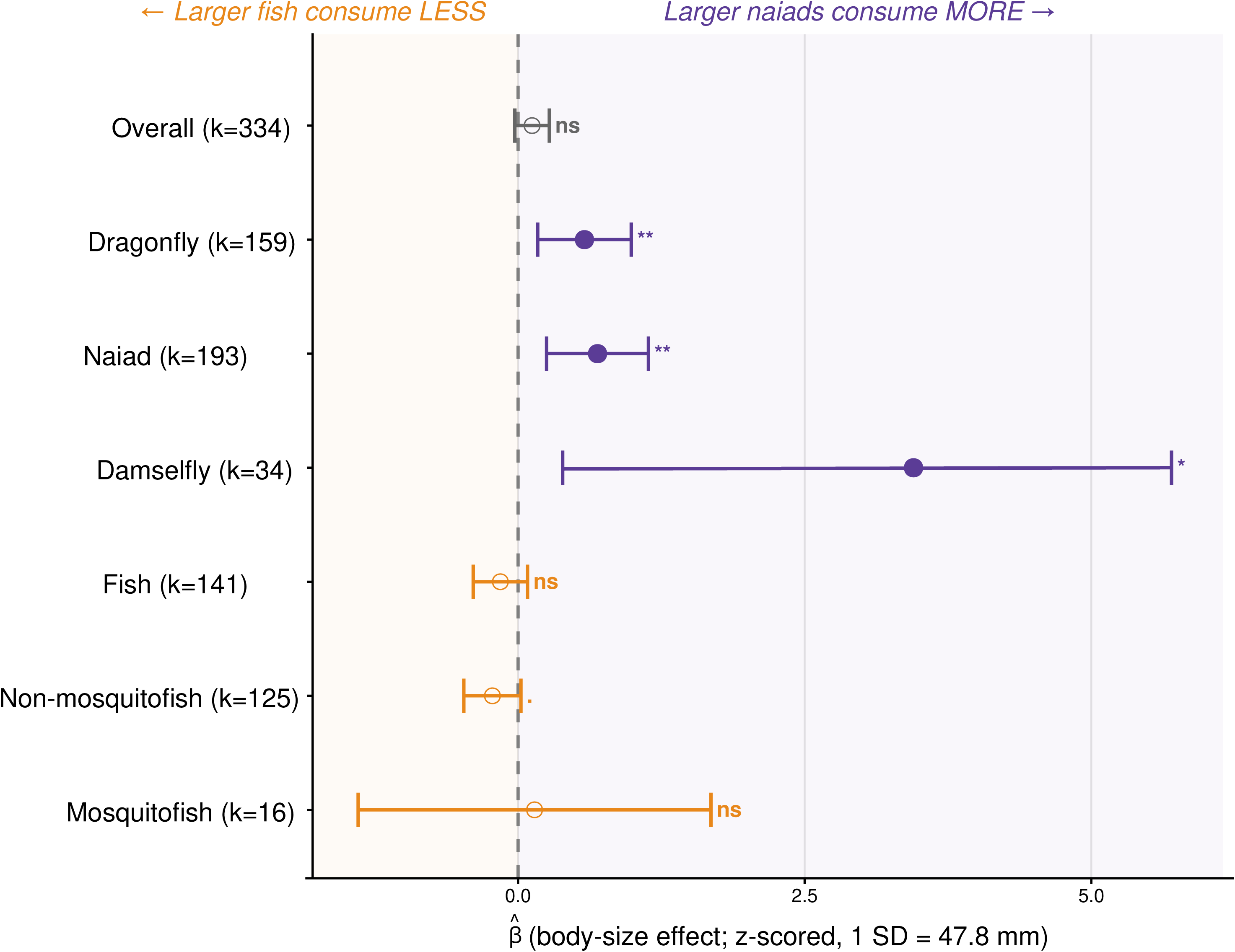
Coefficient plot summarising body-size effects (β, global z-scored units; 1 SD = 47.8 mm) across all predator groups, with 95% CIs. Filled symbols: p < 0.05; open symbols: p ≥ 0.05. Grey-shaded background distinguishes fish groups (negative or flat slopes) from naiad groups (positive slopes) — rendered in greyscale/pattern fill rather than colour per journal formatting. Dragonfly naiads are the only group with a statistically significant positive relationship.

### 3.5 Practical range of biocontrol efficacy

Considering each significant moderator independently, we can estimate the maximum potential practical range of predation across predator–prey combinations. These predictions combine the significant single-trait effects identified above under an additivity assumption, and so provide an approximate, upper-bound estimate rather than a tested joint model. Because body size further predicts CR in naiads but not in fish, with mosquitofish being the most efficient guild tested (CR = 3.36), we argue that fish body size may already sit close to a practical ceiling for predation rate, whereas a sufficiently large dragonfly naiad can exceed even this: the most favourable predicted combination in our dataset (a large dragonfly naiad against *Aedes* early instars in a 1-hour assay) yields an estimated CR of 10.1 larvae predator □¹ h □¹. Where information on local naiad body size is unavailable, mosquitofish remains the best-supported single choice among the guilds we tested. At the other extreme, the least efficient combination (non-mosquitofish against *Culex* late instars in a 24-hour assay) yields CR ≈ 0.08 — a 127-fold range that illustrates how strongly predator and prey traits jointly determine realistic biocontrol expectations; these figures should be interpreted as approximate, trait-based bounds rather than site-specific predictions.

## 4. Discussion

Our meta-analysis provides a comprehensive quantitative synthesis of predator-mediated biological control of mosquito larvae, integrating predator taxonomic identity, developmental stage, dietary guild, body size, and exposure duration as joint predictors of predation efficacy within a single analytical framework. Expressing predation efficiency as a standardised consumption rate (CR, larvae predator □¹ h □¹; defined in Section 2) allowed direct comparability across studies differing substantially in experimental duration. Five main results emerged, one from each of the principal analyses presented above. First, predator identity mattered far more at fine taxonomic resolution than at the broad fish-versus-naiad level, with mosquitofish and dragonfly naiads achieving comparably high consumption rates while non-mosquitofish fish lagged well behind both. Second, exposure duration was a strong, general modulator of consumption rate across all predator groups, with short assays consistently overestimating sustained efficacy. Third, dietary guild was the clearest trait-based predictor of efficacy in fish, with insectivorous species consistently outperforming omnivorous and herbivorous species. Fourth, body size was a strong positive predictor of consumption rate in naiads but not in fish, pointing to fundamentally different foraging constraints between the two groups. Fifth, combining these traits across realistic predator–prey scenarios revealed a wide practical range of biocontrol outcomes, underscoring that no single predator or trait combination is universally optimal. Together, these results move the evidence base for mosquito biocontrol from broad taxonomic assumptions toward an explicit, trait-based framework for predator selection.

### 4.1 Dragonfly naiads rival mosquitofish as mosquito predators

Our results challenge the routine use of broad taxonomic categories in biocontrol assessments. Among the four specific predator groups, mosquitofish and dragonfly naiads had the highest, statistically indistinguishable consumption rates, while non-mosquitofish fish were by far the least efficient group. This contradicts the implicit assumption that fish are generally more effective biocontrol agents than naiads (Onen et al., 2024; Cuthbert et al., 2020; Walshe et al., 2017). The pattern instead points to dietary specialisation, rather than taxonomic identity, as the operative trait: mosquitofish are highly insectivorous surface feeders, and dragonfly naiads are opportunistic, non-specialised macroinvertebrate predators, whereas most non-mosquitofish species in our dataset feed on a broader, less insect-dominated diet. Framed this way, predator performance tracks functional feeding ecology, not phylogeny, and the same trait-matching logic that favours mosquitofish over generic “fish” also favours dragonfly naiads over generic “naiads”.

We also found that fish consumption rates were strongly modulated by the genus of the mosquito, with fish highly efficient against *Aedes* but considerably less so against *Culex*, whereas naiads showed consistent rates across all three genera. This genus-specificity in fish plausibly reflects differences in larval behaviour and habitat use: *Aedes* larvae typically occupy small, structurally simple container habitats where visually orienting fish predators (if any) forage unimpeded, whereas *Culex* larvae are more often associated with vegetated, organically enriched habitats that provide refuge from fish; *Anopheles*, which forages near the water surface, fell at intermediate rates. This differential vulnerability is directly relevant to vector control, since *Culex* is a principal vector of West Nile virus and lymphatic filariasis (Akter et al., 2023): the comparatively poor performance of fish against this genus, together with the consistent performance of naiads across genera, suggests naiads offer more reliable, genus-independent control in mixed mosquito assemblages. The consistent cross-stage and cross-genus efficacy of naiads likely reflects their ambush-foraging strategy: rather than actively pursuing prey, naiads remain stationary and capture larvae that enter their strike range, so their consumption rate depends primarily on encounter probability — a function of naiad density and habitat structure — rather than on active search effort or the specific escape behaviour of a given mosquito genus or larval instar (Russell et al., 2022). This mechanistic difference in foraging mode, rather than any general “naiad advantage”, best explains why naiad performance was more genus- and stage-invariant than fish performance throughout our results.

### 4.2 Diet specialisation predicts predation efficiency: insectivorous fish outperform generalists

Considering dietary specialisation among fish, insectivorous species consumed substantially more larvae per hour than herbivorous species, with omnivorous species intermediate; critically, insectivorous fish also substantially outperformed omnivorous fish in the formal moderator model, indicating that the insectivore advantage is not merely an artefact of comparison against a sparsely represented herbivore guild but a robust, model-supported effect. This gradient likely reflects the suite of morphological and behavioural traits associated with insectivory — visual acuity, strike speed, and gape morphology suited to capturing mobile invertebrate prey (Gerking, 1994) — traits that herbivorous and omnivorous fish, adapted primarily to plant material or detritus, do not share. The two herbivorous-fish studies in our dataset, though too few to model formally, were consistent with this interpretation, showing consumption rates below both other guilds. The bimodal distribution of consumption-rate values among insectivorous fish (Fig. 4) warrants further attention, and future work should test whether prey developmental stage or arena volume is the primary driver of this within-guild heterogeneity. The overall pattern strongly supports trait-based predator selection (Cuthbert et al., 2020; Kalinoski & DeLong, 2016).

The positive relationship between body size and consumption rate in naiads, but not fish, is consistent with fundamentally different constraints operating on the two groups. As sit-and-wait ambush predators, naiads are plausibly gut-capacity (satiation) limited: smaller individuals reach satiation after consuming relatively few larvae, while larger individuals can process substantially more prey before reaching that ceiling, producing the steep positive body-size effect we observed (Weterings et al., 2015). Fish, in contrast, are active foragers whose consumption within a fixed assay period is more plausibly limited by the number of successful pursuit-and-capture events — itself a function of larval evasive behaviour, encounter rate, and handling time — than by digestive capacity; under this interpretation, a larger fish gains little advantage unless its increased size also improves capture success, which our data suggest it does not. This distinction implies that body size is a more reliable, general selection criterion when deploying naiads than when deploying fish, for which dietary guild is the more informative trait (see above); alternative explanations, including a broadening of dietary niche with increasing fish body size (Peters, 1983) or confounds between predator size and arena volume across studies (Uiterwaal & DeLong, 2020), cannot be excluded and warrant testing with standardised arena designs.

A central methodological contribution of this synthesis is the standardised, per-hour consumption-rate metric, which allowed us to directly compare studies with exposure durations ranging from minutes to 24 h that could not otherwise be pooled. This standardisation revealed a strong, consistent negative relationship between exposure time and per-hour consumption rate across all predator groups. Two, non-mutually-exclusive mechanisms likely underlie this pattern: progressive prey depletion, whereby encounter rate necessarily declines as available larvae are consumed over the course of a trial; and predator satiation, whereby consumption slows once gut capacity is approached, regardless of prey availability (Uiterwaal & DeLong, 2020; Englund et al., 2011). A portion of the observed negative slope is also mathematically inherent to the incidence-rate metric itself, and disentangling true biological depletion or satiation from this mathematical component would require factorial designs that orthogonally vary exposure time and prey density. From a practical standpoint, the choice of assay duration should match the management question being asked: brief assays (under about 1 h) are informative about how quickly a predator locates and engages prey, but they systematically overestimate the rate at which a single predator can reduce a standing larval population over a realistic time frame; longer, larger-arena trials that allow genuine prey depletion to occur are more representative of a predator’s likely impact on a natural larval community. We therefore recommend that future efficacy trials adopt multi-duration designs (e.g. 1, 6, and 24 h) to separate instantaneous engagement rate from sustained, depletion-limited biocontrol potential (Walshe et al., 2017); because longer trials also tend to use larger arenas, designs that vary exposure time and arena size independently would further help disentangle prey depletion from encounter-rate artefacts.

Beyond exposure duration, several broader features of the primary literature merit critical assessment. The geographic dominance of Asian studies (roughly two-thirds of the dataset, concentrated in India) and the near-absence of research from Africa, the Americas, and Europe represent a serious mismatch between where this research is conducted and where mosquito-borne disease burden is highest (Dambach et al., 2020). Taxonomically, our synthesis was similarly constrained: naiad data derive predominantly from dragonflies, with damselfly estimates based on comparatively few, uncertain effect sizes, and fish dietary-guild comparisons were limited by a near-absence of herbivorous species. Methodologically, the near-total reliance on laboratory or semi-field assays is a related concern: under natural field conditions, habitat complexity, alternative prey availability, and community-level interactions — including non-consumptive effects of predator cues on larval behaviour (Russell et al., 2022) — will modify the consumption rates estimated here (Alto et al., 2005; Dhanker et al., 2014; Kroeger et al., 2013). Addressing these gaps should be a priority for future research: geographically targeted studies in under-represented, high-burden regions; expanded testing of damselfly naiads and herbivorous fish guilds; and field or semi-field validation of the trait-based predictions derived from this synthesis.

The multilevel prediction interval for consumption rate spanned three orders of magnitude, indicating that the pooled mean is not a useful predictor for any specific application context. That the omnibus model nonetheless explained a substantial share of between-study variance (Section 3) shows, however, that this heterogeneity is far from random: it is structured predominantly by the ecological traits we quantified, which is reassuring for evidence-based deployment even when the overall average is uninformative (Romero & Srivastava, 2010).

Three evidence-based recommendations emerge from synthesising the results of our meta-analysis. First, native odonate naiads — and dragonfly naiads in particular — represent a preferred biocontrol option over most fish in terms of consumption rate, combining per-hour rates comparable to mosquitofish with consistent performance across mosquito genera and larval stages, a statistically robust positive body-size effect, and, unlike the invasive *Gambusia* species conventionally deployed for mosquito control, no documented history of range expansion or ecological harm following introduction (Pyke, 2008); this makes dragonfly naiads a lower-risk choice in ecologically sensitive settings, particularly the small, container-type habitats favoured by *Aedes*, where regulatory or conservation concerns often preclude *Gambusia* release. Second, for fish predators, dietary guild rather than species identity is the primary predictor of biocontrol potential, with insectivorous species consistently outperforming omnivorous species and herbivorous species offering essentially no useful predation; because guild classification is readily available from FishBase (Froese & Pauly, 2025), it should be a primary, low-cost screening criterion when selecting fish biocontrol agents. Third, efficacy estimates are highly sensitive to assay duration: short assays substantially overestimate sustained, 24-hour consumption rates, and this overestimation is most pronounced in non-mosquitofish fish, so multi-duration designs are a prerequisite for meaningful cross-study comparison (Walshe et al., 2017; Uiterwaal & DeLong, 2020). Given these trait-based patterns, the results reported here can themselves inform an initial, quantitative prediction of expected consumption rates for a given predator, mosquito genus, and larval-stage combination; local pilot testing remains most valuable where key covariates absent from our dataset — water turbidity, temperature, or structural habitat complexity, for example — are expected to strongly modify predation, rather than as a universal prerequisite for applying these findings. Taken together, these three recommendations share a common thread: no single predator type is universally superior, and effective biocontrol requires systematic matching of predator functional traits — body size, foraging mode, dietary guild — to the specific mosquito assemblage, larval-stage distribution, and habitat structure of the target site.

## Acknowledgements

[To be completed.]

## Data Availability Statement

Data and analysis code supporting the results of this study will be deposited at the Zenodo Digital Repository upon acceptance (DOI to be assigned). They are available upon request (contact).

## Author Contributions

Nuñez JD: Conceptualisation, Data curation, Formal analysis, Methodology, Writing – original draft. Jolles JW: Conceptualisation, Supervision, Writing – review & editing. Bartumeus F: Conceptualisation, Funding acquisition, Supervision, Writing – review & editing.

