## Supplementary Material for "Mosquito population control by aquatic predators: a global meta-analysis of predation efficacy by fish and odonate naiads"

*Journal of Animal Ecology*

##### Contents


- Table S1 – Data extraction template variables
- Table S2 – Bayesian model convergence diagnostics
- Table S3 – Multilevel heterogeneity statistics
- Table S4 – Publication bias analyses
- Table S5 – Leave-one-out influence diagnostics (summary)
- Table S6 – Cluster-robust standard errors (Predator type model)
- Table S7 – Predator type × Mosquito genus full coefficients
- Table S8 – Predator type × Larval stage full coefficients
- Table S9 – Specific predator × Larval stage full coefficients
- Table S10 – Within-group body size slopes
- Table S11 – Exposure time practical consequences by predator group
- Figure S1 – PRISMA flow chart for study inclusion
- Figure S2 – Funnel plot

Table S1: **Table S1.** Variables captured in the standardised data extraction template. Variables are grouped into six categories.

| **Category** | **Variable** | **Description** |
|---|---|---|
| Study characteristics | StudyID | Unique study identifier |
| | Country | Country of study |
| | Latitude/Longitude | Geographic coordinates |
| | Year | Publication year |
| Predator attributes | Predator species | Taxonomic identity |
| | Predator family | Family-level classification |
| | Body size (mm) | Predator total length |
| | Number of predators | Predators per experimental unit |
| | Feeding mode | Insectivorous / Omnivorous / Herbivorous |
| Prey characteristics | Mosquito genus | *Aedes* / *Anopheles* / *Culex* |
| | Larval instar | Early (I–II) / Late (III–IV) |
| | Number offered | Initial larvae per replicate |
| Experimental design | Duration | Exposure time (hours) |
| | Replicates | Number of independent experimental replicates (trials) |
| Predation metrics | Larvae consumed | Mean larvae consumed per unit |
| | Larvae remaining | Mean larvae surviving per unit |
| Statistics | SD / SE | Standard deviation or error |
| | Sample size | Number of replicates |

Table S2: **Table S2.** Bayesian model convergence diagnostics and comparison with frequentist estimates. Models fitted using brms 2.20.4 with four MCMC chains (6000 iterations, 2000 warm-up), weakly informative priors Normal(0, 1) for coefficients and half-$t(3)$ for variance components. $\hat{R}$: Gelman–Rubin diagnostic ($< 1.01$ indicates convergence). ESS: effective sample size. CrI: credible interval.

| **Parameter** | **Freq. logIR** | **Bayes logIR** | **95 % CrI** | $\hat{R}$ | **ESS** | **P(dir.)** |
|---|---|---|---|---|---|---|
| Overall intercept | 0.965 | 0.903 | [0.589, 1.220] | 1.01 | 450 | 1.00 |
| Fish (intercept) | 0.829 | 0.724 | [0.326, 1.120] | 1.00 | 492 | 1.00 |
| Naid contrast | +0.273 | 0.340 | [0.056, 0.623] | 1.01 | 1260 | 0.99 |
| Early instar | 1.302 | 1.210 | [0.835, 1.580] | 1.02 | 454 | 1.00 |
| Late (contrast) | –0.374 | –0.374 | [–0.84, –0.43] | ≤1.007 | > 400 | 0.35 |
| Insectivore | +1.428 | +1.45 | [+0.51, +2.39] | ≤1.007 | > 400 | 0.81 |
| Herbivore | +0.327 | +0.24 | [–0.65, +1.12] | ≤1.007 | > 400 | 0.70 |
| Naid×Late | +0.212 | +0.15 | [–0.37, +0.63] | ≤1.007 | > 400 | 0.72 |

Table S3: **Table S3.** Multilevel heterogeneity statistics for the intercept-only model ($k = 755$; $n_{\text{st}} = 59$ studies), quantifying how much of the total variance in consumption rate reflects genuine between- and within-study heterogeneity rather than sampling error alone. $QE$ is Cochran's $Q$ test for residual heterogeneity; its highly significant value, together with $I^2_{\text{total}} = 99.1\,\%$, indicates that differences in consumption rate across effect sizes are overwhelmingly due to true heterogeneity, most of which occurs between studies ($I^2_{\text{between}} = 77.5\,\%$) rather than within them ($I^2_{\text{within}} = 21.6\,\%$). Estimated from variance components following Nakagawa et al. (2015, *Methods in Ecology and Evolution*).

| **Parameter** | **Value** |
|---|---|
| $\sigma^2_{\text{between-study}}$ (StudyID) | 1.628 |
| $\sigma^2_{\text{within-study}}$ (SubID) | 0.453 |
| Typical sampling variance ($\bar{v}_i$) | 0.019 |
| $I^2_{\text{total}}$ | 99.1 % |
| $I^2_{\text{between-study}}$ | 77.5 % |
| $I^2_{\text{within-study}}$ | 21.6 % |
| $QE$ (df = 754) | > 50000 |
| $p$-value ($QE$) | < 0.001 |
| 95 % Prediction interval (IRLN) | [−1.88, +3.81] |
| 95 % Prediction interval (IR) | [0.15, 45.15] |

Table S4: **Table S4.** Publication bias analyses, testing whether small, low-precision studies systematically report larger effects than large, high-precision studies. All analyses based on the overall intercept model ($k = 755$). Egger's test and Begg's $\tau$ both indicate significant funnel-plot asymmetry, but under the extreme heterogeneity of this dataset such asymmetry is expected even without genuine publication bias; PET-PEESE, which is robust to this issue, gives a corrected estimate (IR = 3.17) reasonably close to the uncorrected estimate (IR = 2.62), whereas trim-and-fill's implausible imputation of 195 missing studies (more than three times the 59 observed) confirms its known unreliability here. PET-PEESE: Precision Effect Test – Precision Effect Estimate with Standard Errors (weighted regression of logIR on $\sqrt{v_i}$ or $v_i$ in a multilevel model). Trim-and-fill: single-level `rma` required; multilevel estimate used for comparison. Note: imputing 195 studies far exceeds the 59 observed, likely reflecting the documented failure of trim-and-fill under $I^2 \approx 99\,\%$ (Terrin et al. 2003); PET-PEESE is the preferred correction.

| **Method** | **logIR** | **IR** | **Note** |
|---|---|---|---|
| Original (rma.mv) | 0.965 | 2.62 | — |
| Trim-and-fill (left side) | 1.014 | 2.76 | $k_0 = 0$ imputed |
| Trim-and-fill (right side) | 1.514 | 4.54 | $k_0 = 195$ imputed |
| Egger's intercept (multilevel) | 1.772 | 5.88 | $p < 0.001$ |
| PET (SE as predictor; multilevel) | 1.772 | 5.88 | $p < 0.001$ |
| PEESE ($v_i$ as predictor; multilevel) | 1.155 | 3.17 | $p < 0.001$ |
| Begg's $\tau$ | $\tau = -0.160$, $p < 0.001$ | | |

Table S5: **Table S5.** Leave-one-out influence diagnostics. For each of the 59 studies, the overall intercept model was re-fitted with that study removed. $\delta$: absolute change in overall logIR estimate. $\delta_{\%}$: percentage change. Maximum influence study: P3 (internal study identifier; see Supplementary Data S1 for the full study list; $\delta = 4.44\,\%$ of overall estimate), below the conventional $> 10\,\%$ threshold for influential observations.

| **Metric** | **Value** |
|---|---|
| Number of studies | 59 |
| Most influential study | P3 |
| Maximum $\lvert\delta\rvert$ (logIR) | 0.043 |
| Maximum $\lvert\delta_{\%}\rvert$ | 4.44 % |
| Studies with $\lvert\delta_{\%}\rvert > 10\,\%$ | 0 |
| Conclusion | No influential studies detected |

Table S6: **Table S6.** Cluster-robust standard errors (type CR2; Rodgers & Pustejovsky 2020) for the predator type model, addressing the fact that multiple effect sizes drawn from the same study are not statistically independent and would otherwise understate the true uncertainty of the model-averaged estimates. Standard errors are clustered at the study level to account for non-independence; these robust SEs, rather than the model-based SEs, were used to verify the significance of the main effects reported in Table 1.

| **Parameter** | **Estimate** | **SE (robust)** | $t$ | $p$ |
|---|---|---|---|---|
| Intercept (Fish) | 0.829 | 0.314 | 2.61 | 0.0123 |
| Naid contrast | 0.273 | 0.480 | 0.565 | 0.599 |

Table S7: **Table S7.** Full coefficients for the Predator type × Mosquito genus interaction model ($k = 755$; $QM_{(5)} = 17.88$, $p = 0.003$). Reference levels: Fish × *Aedes*.

| **Parameter** | **Estimate** | **SE** | **95 % CI** | $p$ |
|---|---|---|---|---|
| Intercept (Fish×*Aedes*) | 1.162 | 0.243 | [0.687, 1.638] | < 0.001 |
| Naiad contrast | 0.025 | 0.208 | [−0.383, +0.433] | 0.905 |
| *Anopheles* contrast | −0.288 | 0.222 | [−0.724, +0.148] | 0.195 |
| *Culex* contrast | −0.594 | 0.213 | [−1.012, −0.177] | 0.005 |
| Naid × *Anopheles* | +0.469 | 0.279 | [−0.079, +1.016] | 0.093 |
| Naid × *Culex* | +0.411 | 0.248 | [−0.075, +0.898] | 0.097 |

Interpretation: fish significantly less efficient against *Culex* than *Aedes* ($p = 0.005$); naiads show non-significant trends towards compensating this genus effect (both $p \approx 0.10$).

Table S8: **Table S8.** Full coefficients for the Predator type × Larval stage interaction model ($k = 739$; $QM_{(3)} = 35.82$, $p < 0.001$). Reference levels: Fish × Early instars. IR values back-transformed from model estimates.

| **Parameter** | **Estimate** | **SE** | **95 % CI** | $p$ | **IR** |
|---|---|---|---|---|---|
| Intercept (Fish:Early) | 1.473 | 0.223 | [1.035, 1.911] | < 0.001 | 4.36 |
| Naid contrast | −0.222 | 0.180 | [−0.575, +0.131] | 0.218 | — |
| Late contrast (fish) | −0.547 | 0.146 | [−0.834, −0.261] | < 0.001 | — |
| Naid×Late | +0.212 | 0.164 | [−0.109, +0.534] | 0.195 | — |
| Derived group IRs: | | | | | |
| Fish:Early | — | — | — | — | 4.36 |
| Fish:Late | — | — | — | — | 2.52 |
| Naid:Early | — | — | — | — | 3.49 |
| Naid:Late | — | — | — | — | 2.50 |
| Stage-dependent decline: Fish 42 %; Naid 28 % (interaction non-significant, $p = 0.195$). | | | | | |

Table S9: **Table S9.** Full coefficients for the Specific predator × Larval stage interaction model ($k = 739$; $QM_{(7)} = 63.74$, $p < 0.001$). Reference level: Mosquitofish × Early instars.

| **Parameter** | **Estimate** | **SE** | **95 % CI** | $p$ | **Sig.** |
|---|---|---|---|---|---|
| Intercept (Mosquitofish:Early) | 2.614 | 0.487 | [1.661, 3.568] | < 0.001 | *** |
| Non-mosquitofish contrast | −1.274 | 0.461 | [−2.177, −0.370] | 0.006 | ** |
| Dragonfly contrast | −1.307 | 0.476 | [−2.240, −0.375] | 0.006 | ** |
| Damselfly contrast | −1.539 | 0.495 | [−2.508, −0.569] | 0.002 | ** |
| Late (Mosquitofish) | −1.420 | 0.471 | [−2.343, −0.497] | 0.003 | ** |
| Non-mosquitofish×Late | +0.863 | 0.483 | [−0.082, +1.809] | 0.074 | . |
| Dragonfly×Late | +1.118 | 0.477 | [+0.183, +2.052] | 0.019 | * |
| Damselfly×Late | +0.904 | 0.504 | [−0.084, +1.891] | 0.073 | . |
| Derived group IRs (Early / Late): | | | | | |
| Mosquitofish | 13.65 / 3.30 (76 % decline) | | | | |
| Non-mosquitofish | 3.82 / 2.19 (43 % decline) | | | | |
| Dragonfly | 3.70 / 2.73 (26 % decline; Dragonfly×Late $p = 0.019$) | | | | |
| Damselfly | 2.93 / 1.75 (40 % decline) | | | | |
| Note: Mosquitofish:Early has only $k = 5$ from 2 studies; estimate highly uncertain. | | | | | |

Table S10: **Table S10.** Within-group standardised body size slopes. These complement Table 2 of the main text, which reports slopes in global $z$-scored units ($1\,\mathrm{SD} = 47.8\,\mathrm{mm}$). Within-group slopes use the within-group SD as the unit, providing a more biologically interpretable metric. Both groups analysed using separate `rma` models (REML).

| **Group** | **Within-SD (mm)** | $\hat{\beta}_{\text{within}}$ | **95 % CI** | $p$ | **Si** |
|---|---|---|---|---|---|
| Naiad | 11.4 | +0.165 | [+0.059, +0.271] | 0.002 | * |
| Fish | 61.7 | −0.200 | [−0.505, +0.106] | 0.200 | n |

*Biological interpretation:*
Each 11.4-mm increase in naiad body length (one within-naiad SD) corresponds to an 18 % increase i
per-hour consumption ($e^{0.165} - 1 = 0.179$).

Table S11: **Table S11.** Practical consequences of exposure time on per-hour predation rate estimates. Values show the per-hour IR at each duration as a percentage of the IR estimated from a 1-hour assay, based on the negative slope from Table 2. Overestimation factor: the factor by which a 1-hour assay overestimates the per-hour rate of a 24-hour assay.

| **Group** | $\hat{\beta}$ **(h$^{-1}$)** | **2 h (% of 1h)** | **6 h (% of 1h)** | **12 h (% of 1h)** | **24 h (% of 1h)** | **24 h ov f** |
|---|---|---|---|---|---|---|
| Overall | −0.072 | 93 | 70 | 45 | 19 | |
| Naiad | −0.050 | 95 | 78 | 55 | 32 | |
| Fish | −0.102 | 90 | 60 | 35 | 10 | |
| Dragonfly | −0.053 | 95 | 77 | 53 | 29 | |
| Damselfly | −0.038 | 96 | 83 | 64 | 42 | |
| Mosquitofish | −0.057 | 94 | 75 | 51 | 27 | |
| Non-mosquitofish | −0.109 | 90 | 58 | 32 | 8 | |

*Note*: these estimates partly reflect the mathematical structure of the IRLN metric ($T$ in denominator of ln(

### Figures

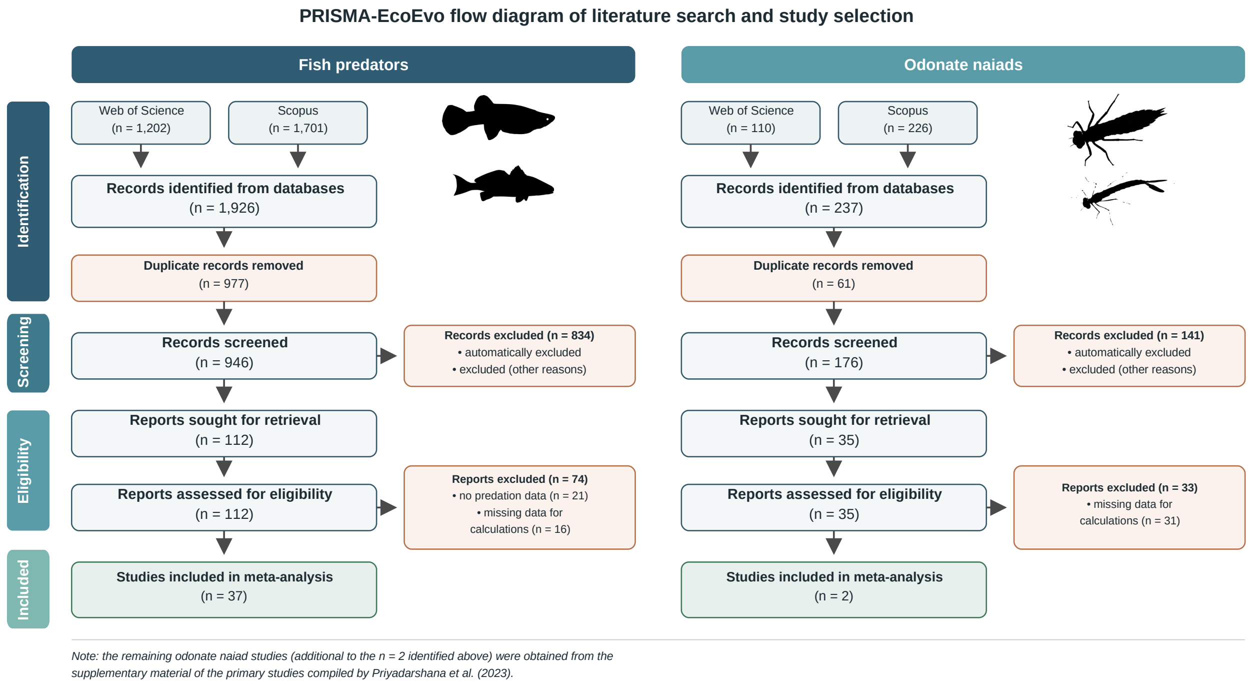


Figure S1: Flowchart of study inclusion for the meta-analysis on predation of mosquito larvae by fish and odonate naiads. Two independent search streams were employed. For fish predators, database searches (Web of Science and Scopus) initially identified 1926 records; after removing duplicates, 946 records were screened; following full-text review, 38 studies were included. For odonate naiads, 33 unique studies were included after integrating the Priyadarshana & Slade (2023) database (supplemented by a targeted update search) and applying our IRLN standardisation. Six studies contributed effect sizes to both predator groups.

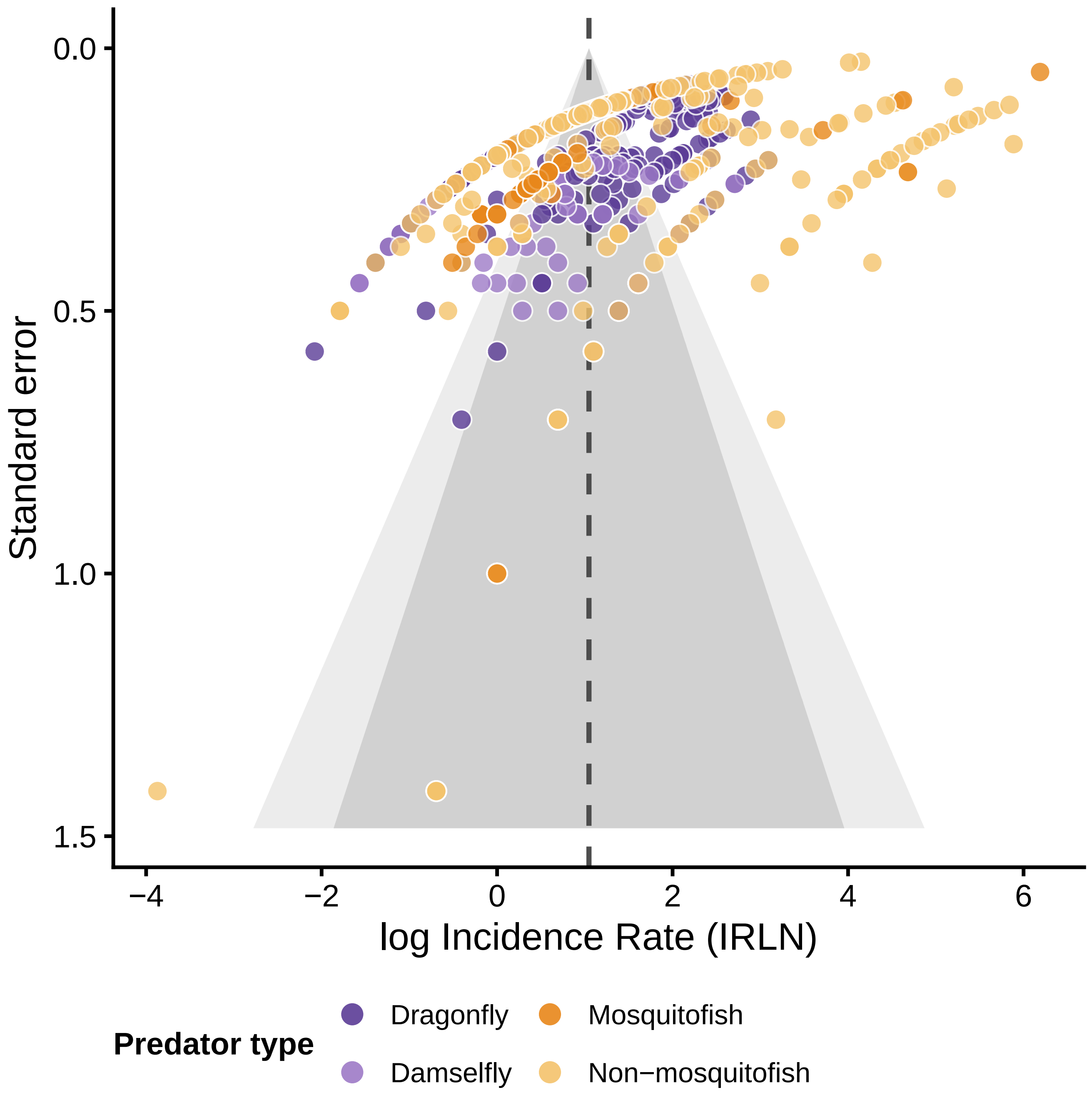


Figure S2: Funnel plot of log incidence rate (IRLN) versus standard error for all 755 effect sizes, coloured by predator group (dragonfly: dark purple; damselfly: light purple; mosquitofish: orange; non-mosquitofish : pale yellow). The dashed vertical line indicates the overall pooled estimate (IRLN = 0.965). Funnel-plot asymmetry (Kendall's $\tau = -0.160$, $p < 0.001$; Egger's intercept = 1.772, $p < 0.001$) indicates publication bias towards larger positive effects; however, under the extreme heterogeneity of this dataset ($I^2 = 99.1\,\%$), asymmetry is expected even in the absence of publication bias (Terrin et al. 2003).

### References (Supplementary)


1. Nakagawa S, et al. Meta-analysis of variation: ecological and evolutionary applications and beyond. *Methods in Ecology and Evolution* 2014;6:143–152.

2. Rodgers MA, Pustejovsky JE. Evaluating meta-analytic methods to detect selective reporting in the presence of dependent effect sizes. *Psychological Methods* 2021;25:719–738.

3. Terrin N, et al. Adjusting for publication bias in the presence of heterogeneity. *Statistics in Medicine* 2003;22:2113–2126.

4. Priyadarshana TS, Slade EM. A meta-analysis reveals that dragonflies and damselflies can provide effective biological control of mosquitoes. *Journal of Animal Ecology* 2023;92:1589–1600.
